# SARS-CoV-2 is transmitted via contact and via the air between ferrets

**DOI:** 10.1101/2020.04.16.044503

**Authors:** Mathilde Richard, Adinda Kok, Dennis de Meulder, Theo M. Bestebroer, Mart M. Lamers, Nisreen M.A. Okba, Martje Fentener van Vlissingen, Barry Rockx, Bart L. Haagmans, Marion P.G. Koopmans, Ron A.M. Fouchier, Sander Herfst

## Abstract

SARS-CoV-2, a coronavirus that newly emerged in China in late 2019 ^1,2^ and spread rapidly worldwide, caused the first witnessed pandemic sparked by a coronavirus. As the pandemic progresses, information about the modes of transmission of SARS-CoV-2 among humans is critical to apply appropriate infection control measures and to slow its spread. Here we show that SARS-CoV-2 is transmitted efficiently via direct contact and via the air (via respiratory droplets and/or aerosols) between ferrets. Intranasal inoculation of donor ferrets resulted in a productive upper respiratory tract infection and long-term shedding, up to 11 to 19 days post-inoculation. SARS-CoV-2 transmitted to four out of four direct contact ferrets between 1 and 3 days after exposure and via the air to three out of four independent indirect recipient ferrets between 3 and 7 days after exposure. The pattern of virus shedding in the direct contact and indirect recipient ferrets was similar to that of the inoculated ferrets and infectious virus was isolated from all positive animals, showing that ferrets were productively infected via either route. This study provides experimental evidence of robust transmission of SARS-CoV-2 via the air, supporting the implementation of community-level social distancing measures currently applied in many countries in the world and informing decisions on infection control measures in healthcare settings ^3^.

---

In late December 2019, clusters of patients in China presenting with pneumonia of unknown etiology were reported to the World Health Organization (WHO) ^1^. The causative agent was rapidly identified as being a virus from the *Coronaviridae* family, closely related to the severe acute respiratory syndrome coronavirus (SARS-CoV) ^2,4,5^. The SARS-CoV epidemic affected 26 countries and resulted in more than 8000 cases in 2003. The newly emerging coronavirus, named SARS-CoV-2 ^6^, rapidly spread worldwide and was declared pandemic by the WHO on March 11, 2020 ^7^. The first evidence suggesting human-to-human transmission came from the descriptions of clusters among the early cases ^8,9^. Based on epidemiological data from China before measures were taken to control the spread of the virus, the reproductive number R0 (the number of secondary cases directly generated from each case) was estimated to be between 2 and 3 ^10–12^. In order to apply appropriate infection control measures to reduce the R0, the modes of transmission of SARS-CoV-2 need to be elucidated. Respiratory viruses can be transmitted via direct and indirect contact (via fomites), and through the air via respiratory droplets and/or aerosols. Transmission via respiratory droplets (> 5 μm) is mediated by expelled particles that have a propensity to settle quickly and is therefore reliant on close proximity between infected and susceptible individuals, usually within 1 m of the site of expulsion. Transmission via aerosols (< 5 μm) is mediated by expelled particles that are smaller in size than respiratory droplets and can remain suspended in the air for prolonged periods of time, allowing infection of susceptible individuals at a greater distance from the site of expulsion ^13^. Current epidemiological data suggest that SARS-CoV-2 is transmitted primarily via respiratory droplets and contact ^8–10,14,15^, which is used as the basis for mitigation of spread through physical and social distancing measures. However, scientific evidence that SARS-CoV-2 can be efficiently transmitted via the air is weak.

Previous studies have shown that ferrets were susceptible to infection with SARS-CoV ^16–20^, and that SARS-CoV was efficiently transmitted to co-housed ferrets via direct contact ^16^. Here, we used a ferret transmission model to assess whether SARS-CoV-2 spreads through direct contact and/or through the air (via respiratory droplets and/or aerosols). For this purpose, individually housed donor ferrets were inoculated intranasally with a strain of SARS-CoV-2 isolated from a German traveller returning from China. Six hours post-inoculation (hpi), a direct contact ferret was added to each of the cages. The next day, indirect recipient ferrets were placed in adjacent cages, separated from the donor cages by two steel grids, 10 cm apart, allowing viruses to be transmitted only via the air (Supplementary Figure 1). On alternating days to prevent cross-contamination, throat, nasal and rectal swabs were collected from each ferret in the inoculated and direct contact groups and from the indirect recipient group, followed by SARS-CoV-2 detection by RT-qPCR and virus titration. Ferrets were productively infected by SARS-CoV-2 upon intranasal inoculation, as demonstrated by the robust and long-term virus shedding from the donor ferrets (Figure 1, Supplementary Figure 2). SARS-CoV-2 RNA levels peaked at 3 days post-inoculation (dpi) and were detected up to 11 dpi in two animals and up to 15 and 19 dpi in the other two animals (Figure 1, Supplementary Figure 2). SARS-CoV-2 was transmitted to direct contact ferrets in four out of four independent experiments between 1 and 3 days post-exposure (dpe) and viral RNA was detected up to 13 to 15 days (i.e. 13 to 17 dpe) (Figure 1, Supplementary Figure 2). Interestingly, SARS-CoV-2 was also transmitted via the air to three out of four indirect recipient ferrets. SARS-CoV-2 RNA was detected from 3 to 7 dpe onwards these indirect recipient ferrets and for 13 to 19 days (Figure 1, Supplementary Figure 2).

**Figure 1.**
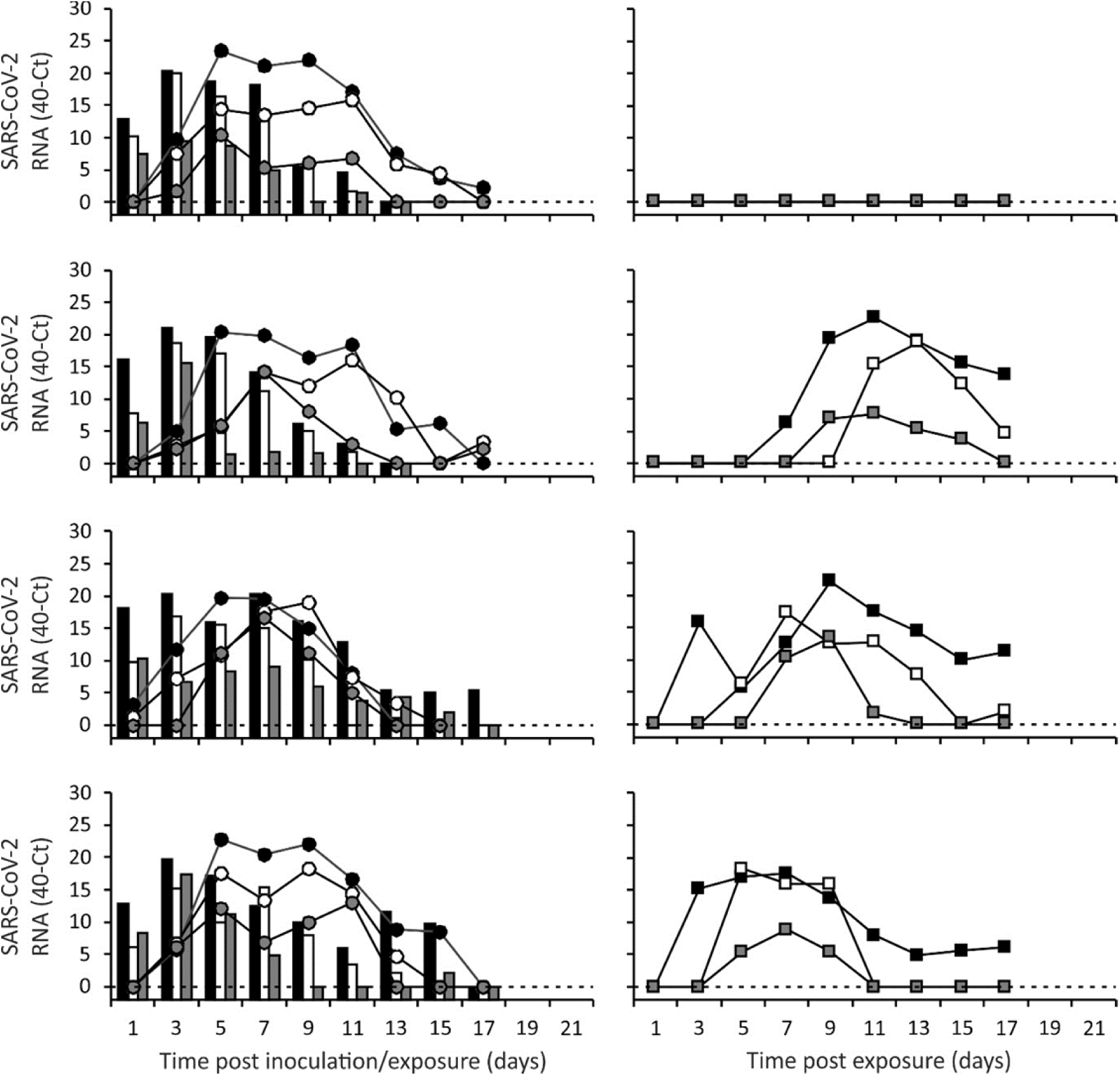
SARS-CoV-2 shedding in ferrets in the transmission experiment. SARS-CoV-2 viral RNA was detected by RT-qPCR in throat (black), nasal (white) and rectal (grey) swabs collected from inoculated donor ferrets (bars; left panels), direct contact ferrets (circles; left panels) and indirect recipient ferrets housed in separate cages (squares; right panels). Swabs were collected from each ferret every other day until no viral RNA was detected in any of the three swabs. The dotted line indicates the detection limit.

**Figure 2.**
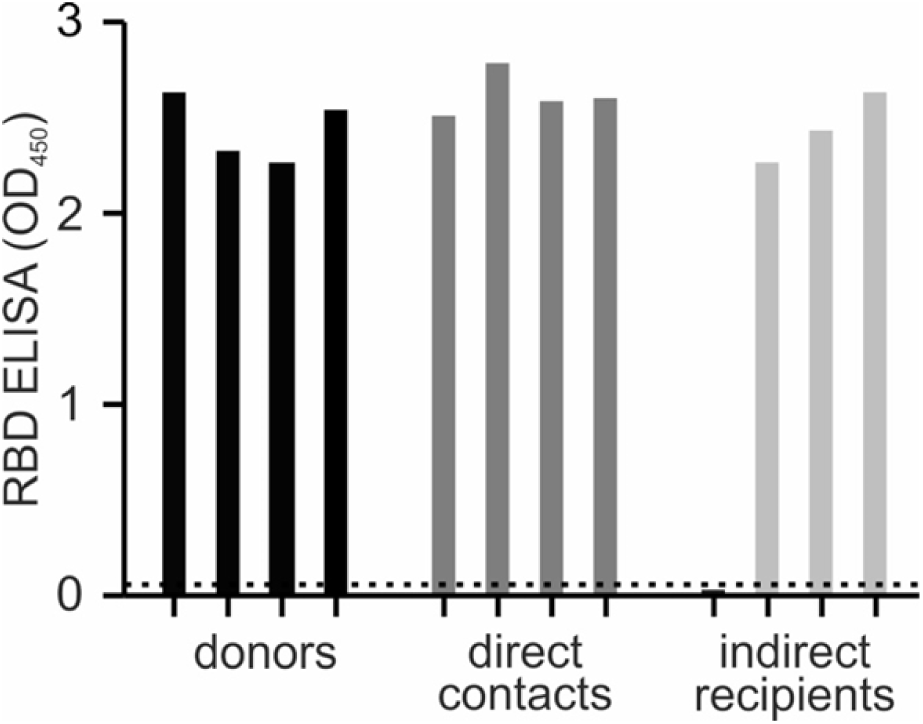
Antibody responses in donor, direct contact and indirect recipient ferrets at 21 dpi/dpe. Sera were collected from the donor, direct contact and indirect recipient ferrets at 21 dpi/dpe and IgG responses were assessed using a SARS-CoV-2 receptor binding domain (RBD) ELISA. The dotted line indicates the background of the assay.

Whereas donor ferrets were inoculated with a high virus dose, direct contact and indirect recipient ferrets are likely to have received a low infectious dose via direct contact or via the air. In spite of this, the pattern of virus shedding from the direct contact and indirect recipient ferrets was similar to that of the inoculated donor ferrets, both in terms of duration and SARS-CoV-2 RNA levels, corroborating robust replication of SARS-CoV-2 upon transmission via direct contact and via the air, independent of the infectious dose. In general, higher SARS-CoV-2 RNA levels were detected in the throat swabs as compared to the nasal swabs. SARS-CoV-2 RNA levels in the rectal swabs were overall the lowest. From each SARS-CoV-2 RNA positive animal, infectious virus was isolated in VeroE6 cells from throat and nasal swabs for at least two consecutive days (Supplementary Table 1). In contrast, no infectious virus was isolated from the rectal swabs. Infectious virus titers ranged from 10^0,75^ to 10^2,75^ TCID_50_/ml (median tissue culture infectious dose per ml) in the donor ferrets, from 10^0,75^ to 10^3,5^ TCID_50_/ml in the direct contact ferrets and from 10^0,75^ to 10^4,25^ TCID_50_/ml in the indirect recipient ferrets. All SARS-CoV-2 positive ferrets seroconverted 21 dpi/dpe, and the antibody levels were similar in donor, direct contact and indirect recipient ferrets (Figure 2). The indirect recipient ferret, in which no SARS-CoV-2 was detected, did not seroconvert as expected.

SARS-CoV-2 transmission in experimental animal models has recently also been described by others. SARS-CoV-2 direct contact transmission between ferrets ^21^ and hamsters ^22^ was reported, with similar efficiency as observed in our study. In addition, SARS-CoV-2 was also found to be transmitted via the air in two out of six ferrets ^21^, and in two out of six cats ^23^. However, only low levels of SARS-CoV-2 RNA were detected in nasal washes and feces of the indirect recipient ferrets, and no infectious virus was isolated ^21^. Furthermore, virus shedding was shorter as compared to the donor animals and only one out of the two SARS-CoV-2 RNA positive indirect recipient ferrets seroconverted. Similarly, the transmission via the air between cats was not efficient. SARS-CoV-2 RNA was detected in the feces and tissues of one cat at 3 and 11 dpi respectively and in nasal washes of another cat, but no infectious virus was isolated. Both SARS-CoV-2 RNA positive indirect recipient cats seroconverted. In contrast, the present study showed that SARS-CoV-2 was efficiently transmitted via the air between ferrets, as demonstrated by long-term virus shedding and the presence of infectious virus in the indirect recipient animals, which is comparable to the transmissibility of pandemic influenza viruses in the ferret model ^24^.

To date, there is no evidence of fecal-oral transmission of SARS-CoV-2 in humans. However, the prolonged detection of RNA in consecutive stool samples ^25^ and the environmental contamination of sanitary equipment ^26^ may suggest that the fecal-oral route could be a potential route of transmission of SARS-CoV-2. Here, no infectious virus was retrieved from any of the rectal swabs. Despite this, it cannot be fully excluded that SARS-CoV-2 was also transmitted from donors to direct contact ferrets partly via the fecal-oral route. In the study by Kim *et al.*, ferret fecal material was used to inoculate ferrets, resulting in a productive infection, indicating that infectious SARS-CoV-2 was shed in fecal specimens ^21^.

Our experimental system does not allow to assess whether SARS-CoV-2 was transmitted via the air through respiratory droplets, aerosols or both, as donor and indirect recipient ferret cages are placed only 10 cm apart from each other. In a recent study, SARS-CoV-2 remained infectious in aerosols for at least 3h after aerosolization at high titers in a rotating drum, comparable to SARS-CoV ^27^. Although it is informative to compare the stability of different respiratory viruses in the air, our study provides the additional information that infectious SARS-CoV-2 particles can actually be expelled in the air and subsequently infect recipients. In two other studies, the presence of SARS-CoV-2 in air samples collected in hospital settings was investigated. However, no SARS-CoV-2 RNA was detected in the air sampled in three isolation rooms ^26^, or 10 cm from a symptomatic patient who was breathing, coughing or speaking ^28^. Nevertheless, RNA was detected on the air exhaust outlet of one of the isolation rooms in the first study, suggesting that virus-laden droplets may be displaced by airflows ^26^.

Here we provide the first experimental evidence that SARS-CoV-2 can be transmitted efficiently via the air between ferrets, resulting in a productive infection and the detection of infectious virus in indirect recipients, as a model for human-to-human transmission. Although additional experiments on the relative contribution of respiratory droplets and aerosols to the transmission of SARS-CoV-2 are warranted, the results of this study corroborate the WHO recommendations about transmission precautions in health care settings and the social distancing measures implemented in many countries around the globe to mitigate the spread ^3^. The ferret transmission model will also be useful to understand transmission dynamics and the molecular basis of the transmissibility of SARS-Cov-2 and other betacoronaviruses, which, in the context of the current SARS-CoV-2 pandemic and future pandemic threats, is clearly of utmost importance.

## Methods

### Virus and cells

SARS-CoV-2 (isolate BetaCoV/Munich/BavPat1/2020; kindly provided by Prof. Dr. C. Drosten) was propagated to passage 3 on VeroE6 cells (ATCC) in Opti-MEM I (1X) + GlutaMAX (Gibco), supplemented with penicillin (10,000 IU mL^−1^, Lonza) and streptomycin (10,000 IU mL ^−1^, Lonza) at 37°C in a humidified CO2 incubator. VeroE6 cells were inoculated at an moi of 0.01. Supernatant was harvested 72 hpi, cleared by centrifugation and stored at –80°C. VeroE6 cells were maintained in DMEM (Gibco) supplemented with 10% foetal calf serum (Greiner), 2 mM of L-glutamine (Gibco), 10 mM Hepes (Lonza), 1.5 mg ml−^1^ sodium bicarbonate (NaHCO_3_, Lonza), penicillin (10,000 IU/mL) and streptomycin (10,000 IU/mL) at 37°C in a humidified CO_2_ incubator. All work was performed in a Class II Biosafety Cabinet under BSL-3 conditions at the Erasmus Medical Center.

### Ferret transmission experiment

All relevant ethical regulations for animal testing have been complied with. Animals were housed and experiments were performed in strict compliance with the Dutch legislation for the protection of animals used for scientific purposes (2014, implementing EU Directive 2010/63). Influenza virus, SARS-CoV-2 and Aleutian Disease Virus seronegative 6 month-old female ferrets (*Mustela putorius furo*), weighing 700–1000 g, were obtained from a commercial breeder (TripleF (USA)). Research was conducted under a project license from the Dutch competent authority (license number AVD1010020174312) and the study protocol was approved by the institutional Animal Welfare Body (Erasmus MC permit number 17-4312-02). Animal welfare was monitored on a daily basis. Virus inoculation of ferrets was performed under anesthesia with a mixture of ketamine/medetomidine (10 and 0.05 mg kg−^1^ respectively) antagonized by atipamezole (0.25 mg kg−^1^). Swabs were taken under light anesthesia using ketamine to minimize animal discomfort.

Four donor ferrets were inoculated intranasally with 6.10^5^ TCID_50_ of SARS-CoV-2 virus diluted in 500 μl of phosphate-buffered saline (PBS) (250 μl instilled dropwise in each nostril) and were housed individually in a cage. Six hpi, direct contact ferrets were placed in the same cage as the donor ferrets. One day later, indirect recipient ferrets were placed in an opposite cage separated by two steel grids, 10 cm apart, to avoid contact transmission (Figure S1). Throat, nasal and rectal swabs were collected from the animals every other day, to prevent cross-contamination, until they were negative for SARS-CoV-2 RNA or maximum for 21 dpi/dpe by determined by real-time RT-qPCR as described below. Swabs were stored at −80 °C in transport medium (Minimum Essential Medium Eagle with Hank's BSS (Lonza), 5 g L−^1^ lactalbumine enzymatic hydrolysate, 10% glycerol (Sigma-Aldrich), 200 U ml−^1^ of penicillin, 200 mg ml−^1^ of streptomycin, 100 U ml−^1^ of polymyxin B sulfate (Sigma-Aldrich), and 250 mg ml−^1^ of gentamicin (Life Technologies)) for end-point titration in VeroE6 cells as described below. Ferrets were euthanized at 21 dpi/dpe by heart puncture under anaesthesia. Blood was collected in serum-separating tubes (Greiner) and processed according to the manufacturer’s instructions. Sera were heated for 1h at 60°C and used for the detection of specific antibodies against SARS-CoV-2 as described below. All animal experiments were performed in class III isolators in a negatively pressurized ABSL3+ facility.

### RNA isolation and RT-qPCR

RNA was isolated using an in-housed developed high-throughput method in a 96-well format. Sixty μl of sample were added to 90 μl of MagNA Pure 96 External Lysis Buffer (Roche). A known concentration of phocine distemper virus (PDV) was added to the sample as internal control for the RNA extraction ^29^. The 150 μl of sample/lysis buffer was added to a well of a 96-well plate containing 50 μl of magnetic beads (AMPure XP, Beckman Coulter). After thorough mixing by pipetting up and down at least 10 times, the plate was incubated for 15 minutes (min) at room temperature. The plate was then placed on a magnetic block (DynaMag™-96 Side Skirted Magnet (ThermoFisher Scientific)) and incubated for 3 min to allow the displacement of the beads towards the side of the magnet. Supernatants were carefully removed without touching the beads and beads were washed three times for 30 seconds (sec) at room temperature with 200 μl/well of 70% ethanol. After the last wash, a 10 μl multi-channel pipet was used to remove residual ethanol. Plates were air-dried for 6 min at room temperature. Plates were removed from the magnetic block and 30 μl of PCR grade water was added to each well and mixed by pipetting up and down 10 times. Plates were incubated for 5 min at room temperature and then placed back on the magnetic block for 2 min to allow separation of the beads. Supernatants were pipetted in a new plate and RNA was kept at 4°C. The RNA was directly used for RT-qPCR using primers and probes targeting the E gene of SARS-CoV-2 as previously described ^30^. The primers and probe for PDV detection were described previously ^29^.

### Virus titrations

Throat, nasal and rectal swabs were titrated in quadruplicates in VeroE6 cells. Briefly, confluent VeroE6 cells were inoculated with 10-fold serial dilutions of sample in Opti-MEM I (1X) + GlutaMAX, supplemented with penicillin (10,000 IU mL^−1^), streptomycin (10,000 IU mL^−1^). At one hpi, the first three dilutions were washed twice with media and fresh media was subsequently added to the whole plate. At six dpi, virus positivity was assessed by reading out cytopathic effects. Infectious virus titers (TCID_50_/ml) were calculated from four replicates of each throat, nasal and rectal swabs and from 24 replicates of the virus stock using the Spearman-Karber method.

### Serology

Sera were tested for SARS-CoV-2 antibodies using a receptor binding domain (RBD) enzyme-linked immunosorbent assay (ELISA) as described previously, with some modifications ^31^. Briefly, ELISA plates were coated overnight with either SARS-CoV-2 RBD. After blocking, sera were added and incubated for 1h at 37°C. Bound antibodies were detected using horseradish peroxidase (HRP)-labelled goat anti-ferret IgG (Abcam) and 3,3',5,5'-Tetramethylbenzidine (TMB, Life Technologies) as a substrate. The absorbance of each sample was measured at 450 nm.

### Data availability

All data are available from the corresponding author (S.H.) on reasonable request. No custom software was used in this study.

## Acknowledgements

We thank Prof. Dr. Christian Drosten (Charité – Universitätsmedizin Berlin) for providing the SARS-CoV-2 isolate used in this study and Drs Rik de Swart and Mathieu Sommers for their help with animal ethics and study approval. This work was supported by European Union's *Horizon 2020* research and innovation program *VetBioNet* (grant agreement No *731014*) and NIH/NIAID (contract number HHSN272201400008C). S.H. was funded in part by an NWO VIDI grant (contract number 91715372).

## Author Contributions

M.R. and S.H. conceived, designed, analysed and performed the work. M.R. and S.H. wrote the manuscript. A.K., D.M., T.B., M.L., N.O. helped with performing the work. M.F.V., B.R., B.H., M.K., R.A.M.F. helped with the design of the work, interpretation of the data and manuscript revision. All authors read and approved the final manuscript.

The authors declare no competing interests.

## Additional information

Supplementary Information is available for this paper.

