## Supplementary Information for "SARS-CoV-2 is transmitted via contact and via the air between ferrets"

**A.**

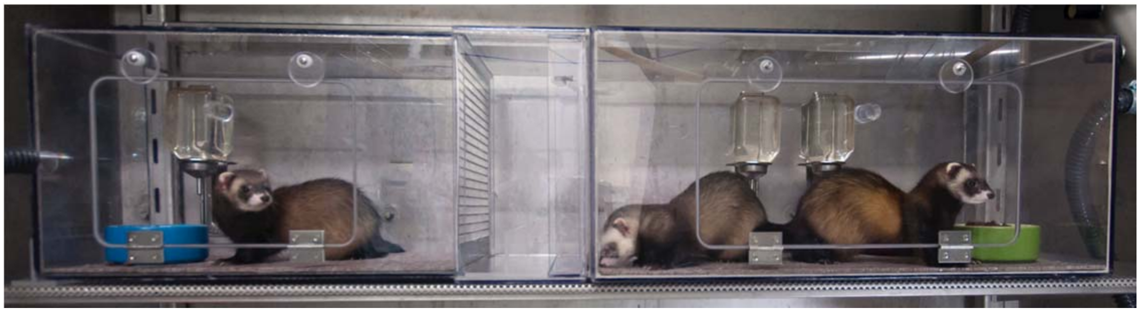

**B.**

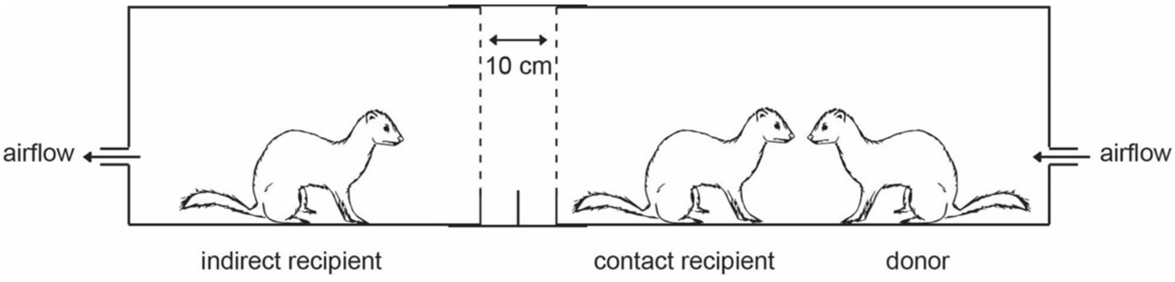


**Supplementary figure 1. The ferret transmission experimental set-up.** Picture (A) and schematic representation (B) of one independent experimental set-up to assess direct contact transmission and indirect transmission via the air. One inoculated donor ferret is housed in a cage (right-hand side of the picture). Six hours later, a direct contact ferret is added to the same cage as the donor ferret. The next day, an indirect recipient ferret is placed in an opposite cage (left-hand side of the picture) separated by two steel grids, 10 cm apart, to avoid contact transmission. The direction of the air flow is indicated by the arrows. The ferret transmission set-ups are placed in class III isolators in a biosafety level 3+ laboratory.

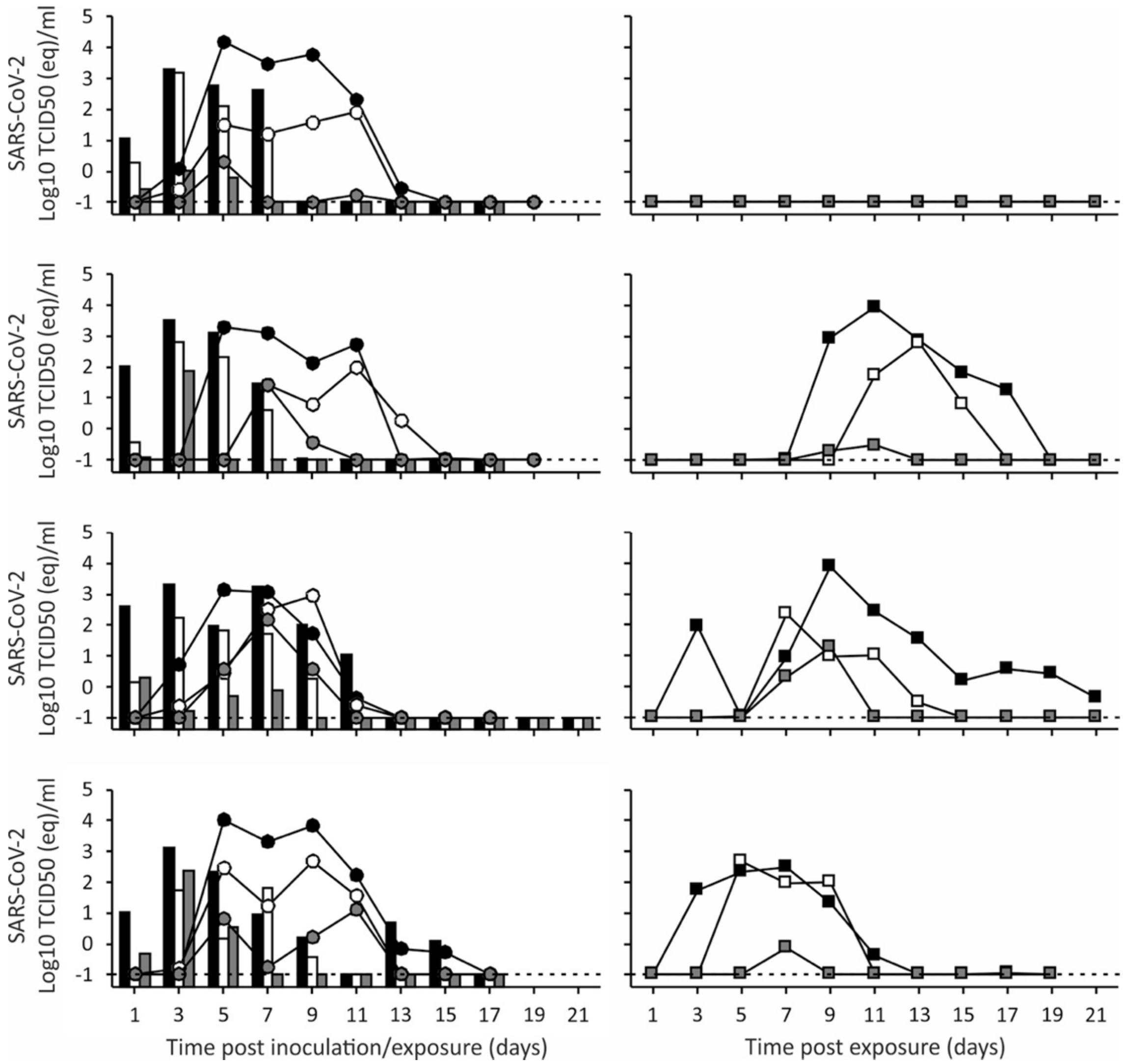


**Supplementary figure 2. SARS-CoV-2 shedding in ferrets in the transmission experiment.** SARS-CoV-2 viral RNA was detected by RT-qPCR in throat (black), nasal (white) and rectal (grey) swabs collected from donor ferrets (bars; left panels), direct contact ferrets (circles; left panels) and indirect recipient ferrets housed in separate cages (squares; right panels). Swabs were collected from each ferret every other day until no viral RNA could be detected in any of the three swabs. $TCID_{50}$ equivalent (eq) were calculated from on a standard curve of serial dilutions of the SARS-CoV-2 viral stock.

**Supplementary Table 1. Detection of SARS-CoV-2 RNA and infectious virus in swabs of ferrets.**

**A** **Donor Ferrets**

| | Nasal swabs | | Throat swabs | | Rectal swabs | |
|---|---|---|---|---|---|---|
| Days post inoculation | RT-qPCR# | Virus culture | RT-qPCR | Virus culture | RT-qPCR | Virus culture |
| 1 | 4/4 | 0/4 | 4/4 | 2/4 (0,75-2,50) | 4/4 | 0/4 |
| 3 | 4/4 | 2/4 (1,75-2,75)* | 4/4 | 4/4 (0,75-2,50) | 4/4 | 0/4 |
| 5 | 4/4 | 0/4 | 4/4 | 2/4 (0,75-1,00) | 4/4 | 0/4 |
| 7 | 4/4 | 1/4 (0,75) | 4/4 | 0/4 | 4/4 | 0/4 |
| 9 | 4/4 | 0/4 | 4/4 | 0/4 | 2/4 | 0/4 |
| 11 | 3/4 | 0/4 | 4/4 | 0/4 | 2/4 | 0/4 |
| 13 | 2/4 | 0/4 | 2/4 | 0/4 | 1/4 | 0/4 |
| 15 | 0/2 | 0/2 | 2/2 | 0/2 | 2/2 | 0/2 |
| 17 | 0/2 | 0/2 | 1/2 | 0/2 | 0/2 | 0/2 |
| 19 | 1/1 | 0/1 | 1/1 | 0/1 | 0/1 | 0/1 |
| 21 | 0/1 | 0/1 | 0/1 | 0/1 | 0/1 | 0/1 |

**B** **Direct contact ferrets**

| | Nasal swabs | | Throat swabs | | Rectal swabs | |
|---|---|---|---|---|---|---|
| Days post inoculation | RT-qPCR | Virus culture | RT-qPCR | Virus culture | RT-qPCR | Virus culture |
| 1 | 1/4 | 0/4 | 1/4 | 0/4 | 0/4 | 0/4 |
| 3 | 4/4 | 0/4 | 4/4 | 0/4 | 3/4 | 0/4 |
| 5 | 4/4 | 2/4 (0,75-1,75) | 4/4 | 4/4 (1,25-3,25) | 4/4 | 0/4 |
| 7 | 4/4 | 3/4 (0,75-1,00) | 4/4 | 4/4 (2,00-3,00) | 4/4 | 0/4 |
| 9 | 4/4 | 2/4 (0,75-1,50) | 4/4 | *4/4* (1,25-3,50) | 4/4 | 0/4 |
| 11 | 4/4 | 0/4 | 4/4 | 1/4 (1) | 4/4 | 0/4 |
| 13 | 4/4 | 0/4 | 3/4 | 0/4 | 0/4 | 0/4 |
| 15 | 1/4 | 0/4 | 3/4 | 0/4 | 0/4 | 0/4 |
| 17 | 1/3 | 0/3 | 1/3 | 0/3 | 1/3 | 0/3 |
| 19 | 0/2 | 0/2 | 0/2 | 0/2 | 0/2 | 0/2 |
| 21 | NT | NT | NT | NT | NT | NT |

**C** **Indirect recipient ferrets**

| | Nasal swabs | | Throat swabs | | Rectal swabs | |
|---|---|---|---|---|---|---|
| Days post inoculation | RT-qPCR | Virus culture | RT-qPCR | Virus culture | RT-qPCR | Virus culture |
| 1 | 0/4 | 0/4 | 0/4 | 0/4 | 0/4 | 0/4 |
| 3 | 0/4 | 0/4 | 2/4 | 0/4 | 0/4 | 0/4 |
| 5 | 2/4 | 1/4 (1,75) | 2/4 | 1/4 (1,00) | 1/4 | 0/4 |
| 7 | 2/4 | 1/4 (1,50) | 3/4 | 1/4 (1,25) | 2/4 | 0/4 |
| 9 | 2/4 | 0/4 | 3/4 | 2/4 (3,00) | 3/4 | 0/4 |
| 11 | 2/4 | 1/4 (1,50) | 3/4 | 1/4 (4,25) | 2/4 | 0/4 |
| 13 | 2/4 | 0/4 | 3/4 | 0/4 | 1/4 | 0/4 |
| 15 | 1/4 | 0/4 | 3/4 | 0/4 | 1/4 | 0/4 |
| 17 | 2/4 | 0/4 | 3/4 | 0/4 | 0/4 | 0/4 |
| 19 | 2/4 | 0/4 | 2/4 | 0/4 | 0/4 | 0/4 |
| 21 | 1/3 | 0/3 | 1/3 | 0/3 | 1/3 | 0/3 |

#Number of animals positive per number of animals tested

*Range of virus titers found in positive samples (in $TCID_{50}$/ml)

NT : not tested
